# Lack of Reinfection in Rhesus Macaques Infected with SARS-CoV-2

**DOI:** 10.1101/2020.03.13.990226

**Authors:** Linlin Bao, Wei Deng, Hong Gao, Chong Xiao, Jiayi Liu, Jing Xue, Qi Lv, Jiangning Liu, Pin Yu, Yanfeng Xu, Feifei Qi, Yajin Qu, Fengdi Li, Zhiguang Xiang, Haisheng Yu, Shuran Gong, Mingya Liu, Guanpeng Wang, Shunyi Wang, Zhiqi Song, Ying Liu, Wenjie Zhao, Yunlin Han, Linna Zhao, Xing Liu, Qiang Wei, Chuan Qin

**Author notes:** These authors contributed equally to this work (Linlin Bao, Wei Deng, Hong Gao, Chong Xiao, Jiayi Liu, Jing Xue, Qi Lv). Correspondence should be addressed to Chuan Qin,.

## Abstract

A global pandemic of Corona Virus Disease 2019 (COVID-19) caused by severe acute respiratory syndrome CoV-2 (SARS-CoV-2) is ongoing spread. It remains unclear whether the convalescing patients have a risk of reinfection. Rhesus macaques were rechallenged with SARS-CoV-2 during an early recovery phase from initial infection characterized by weight loss, interstitial pneumonia and systemic viral dissemination mainly in respiratory and gastrointestinal tracts. The monkeys rechallenged with the identical SARS-CoV-2 strain have failed to produce detectable viral dissemination, clinical manifestations and histopathological changes. A notably enhanced neutralizing antibody response might contribute the protection of rhesus macaques from the reinfection by SARS-CoV-2. Our results indicated that primary SARS-CoV-2 infection protects from subsequent reinfection.

**One Sentence Summary:** Neutralizing antibodies against SARS-CoV-2 might protect rhesus macaques which have undergone an initial infection from reinfection during early recovery days.

---

The Corona Virus Disease 2019 (COVID-19) caused by severe acute respiratory syndrome CoV-2 (SARS-CoV-2), emerged in Wuhan China, has continued to sweep through Europe, America, Asia and more than millions of people have been diagnosed cross the world (*1, 2*). Some patients discharged with undetectable SARS-CoV-2 have been found re-positive during viral detection (*3–5*). Neutralizing antibodies (NAbs) were detected 15 days posterior the onset of COVID-19 (*6–8*). Whether patients have a risk of “relapse” or “reinfection” after recovery from initial infection have aroused the worldwide concern. Therefore, in this study, we used nonhuman primates to track the longitudinally infectious status from primary SARS-CoV-2 infection to reinfection by the same viral strain. Seven adult Chinese-origin rhesus macaques (No M0-M6, 3-5 kg, 3-5-year-old) were modeled for challenge-rechallenge observation. Six monkeys (M1 to M6) were intratracheally challenged with SARS-CoV-2 at 1×10^6^ 50% tissue-culture infectious doses (TCID50). After they underwent mild-to-moderate COIVD-19 and stepped into recovering stage from the primary infection, four monkeys (M3 to M6) were rechallenged intratracheally with the same dose of SARS-CoV-2 strain at 28 days post initial challenge (dpi). Remained two monkeys (M1 and M2) with primary infection were not rechallenged to perform as negative control of rechallenged group. A healthy monkey (M0) was given an initial challenge as model control in the second challenge. The pathological changes with viral-dependent distribution were compared using necropsy specimens between two monkeys undergone challenge-rechallenge (M3 and M5) at 5 days post rechallenge (dpr, 33 dpi) and two monkeys undergone only initial challenge (M0 at 5 dpi and M1 at 7 dpi). Clinical traits including body weight, body temperature, chest X-ray, peripheral blood measurement, nasal/throat/anal swabs, virus distribution, and pathological changes were examined at designated time points (Figure 1). Weight loss ranged from 200 g to 400 g were found in four monkeys undergone initial challenge (4/7) (Figure 2A), and none of monkey’s rectal temperature was observed elevated (0/7) (Figure 2B). Reduced appetite and/or increased respiration were common (6/7) but emerged transiently with a very short time. In regard to viral dissemination, peak viral load (6.5 log10 RNA copies/mL) was detected in nasal swabs and pharyngeal swabs at 3 dpi followed with gradual decline (Figure 2C and 2D). Peak viral load (5 log10 RNA copies/mL) could be detected using anal swabs at 3 dpi followed with linearly declined to undetectable level at 14 dpi (Figure 2E). For all monkeys with the initial challenge, white blood cell (WBC, 3.5-9.5×10^9^/L), lymphocyte counts (LYMP, 1.1-3.4×10^9^/L) and neutrophil counts (NEUT, 1.8-6.4×10^9^/L) fluctuated within normal ranges. Comparing to the baseline, a slight yet significant reduction of WBC and LYMP was observed posterior primary infection (\**p*<0.05, Figure 2F). T lymphocyte subsets including CD4^+^ T cells and CD8^+^ T cells maintained relatively stable during the primary infectious stage (Figure 2G). Specific antibody against SARS-CoV-2 was gradually increased, leading to the concentration significantly higher at 21 dpi compared to that at 3 dpi (\**p*<0.05, Figure 2H). Radiologically, bilateral groundglass opacities were shown, indicating mild-to-moderate interstitial infiltration in monkeys with pneumonia (Represented by M4, Figure 3A). Using necropsy specimens, viral RNA copies were detected in nose (10^6^ to 10^8^ copies/mL), pharynx (10^4^ to 10^6^ copies/mL), lung (10^3^ to 10^7^ copies/mL) and gut (10^4^ to 10^6^ copies/mL) (Figure 3B, upper panel).

**Figure 1.**
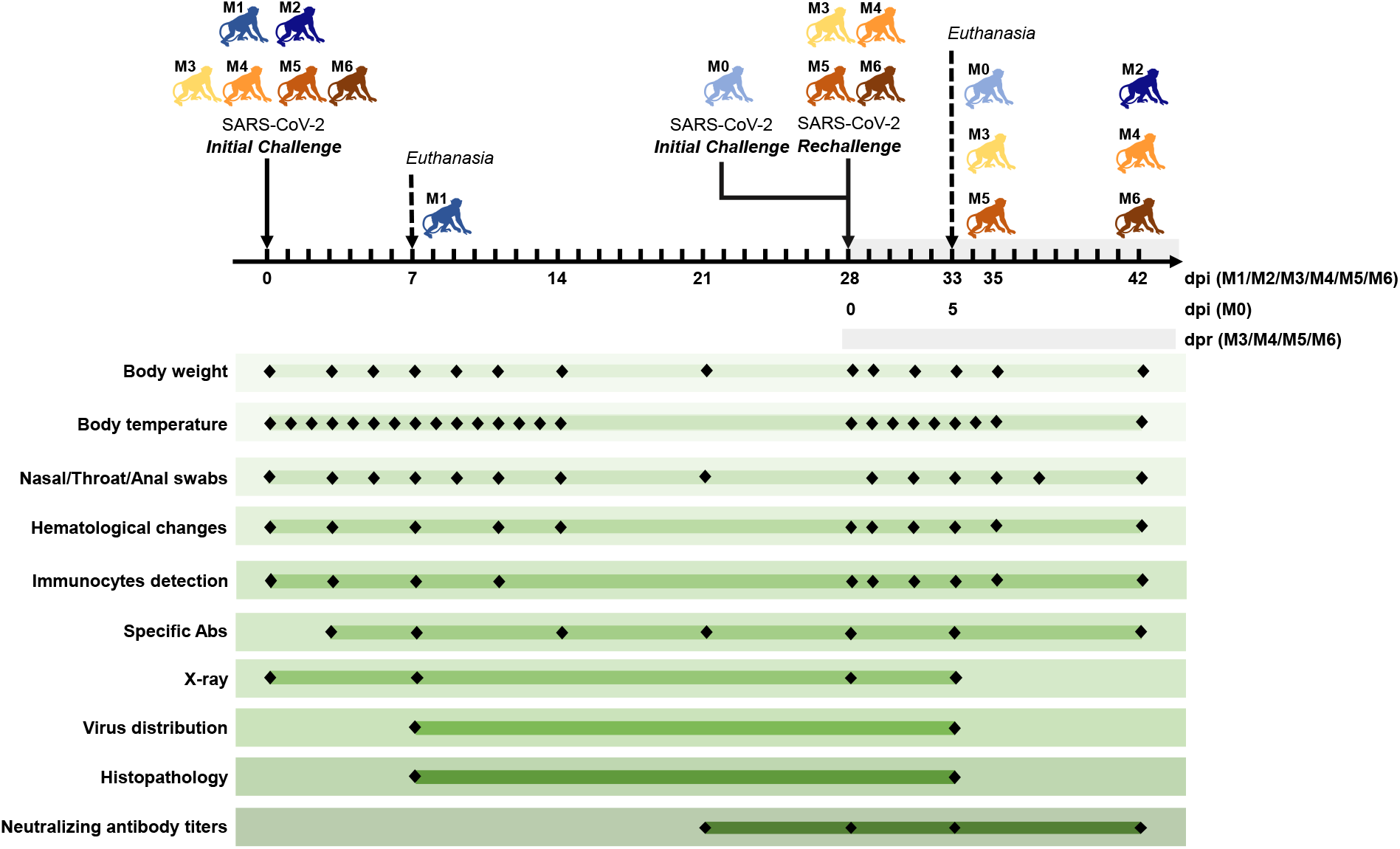
Experimental design and sample collection. Seven adult Chinese-origin rhesus macaques (M0-M6) were enrolled in current study. At the outset of this experiment, six monkeys (M1 to M6) were intratracheally challenged with SARS-CoV-2 at 1×10^6^ TCID50. After all the experimentally infected monkeys were recovery from the primary infection, four infected monkeys (M3 to M6) were intratracheally rechallenged at 28 days post initial challenge (dpi) with the same dose of SARS-CoV-2 strain to ascertain the possibility of reinfection. Meanwhile, uninfected monkey (M0) was also treated with SARS-CoV-2 as the model control in the second challenge, and previously infected monkey (M2) was untreated again and continuously monitored as the control. To compare the virus distribution and histopathological changes between the initially infected monkeys and the reinfected monkeys, two monkeys per group (M0 and M1 in initial infection group, M3 and M5 in reinfection group) were euthanized and necropsied at 5 (M0) or 7 (M1) dpi, 5 (M3 and M5) days post rechallenge (dpr), respectively. Body weight, body temperature, nasal/throat/anal swabs, hematological changes, immunocytes and specific antibody were measured along the timeline at a short interval. Two measurements of virus distribution and histopathology (HE/IHC stain) were carried out at 5 dpi (M0), 7 dpi (M1) and 5 dpr (M3 and M5). Chest X-ray were detected four times and neutralizing antibody titers against SARS-CoV-2 were examined at the indicated time points.

**Figure 2.**
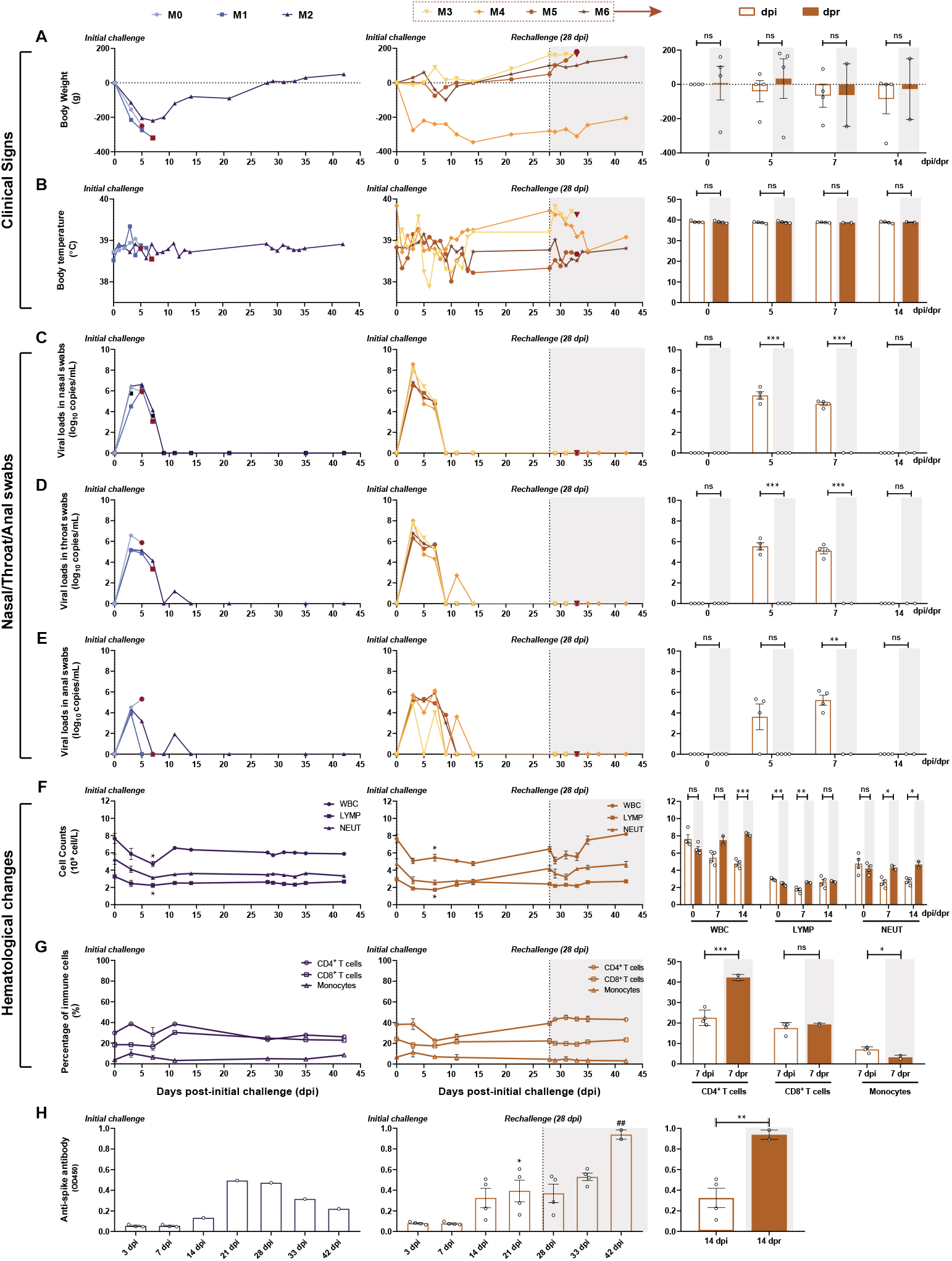
Longitudinally tracking in clinical signs, viral replication, hematological changes and immune response. (**A** and **B**) Clinical signs in each monkey. Monkeys were recorded daily for the changes in body weight and rectal temperature along the timeline after the initial infection followed by the virus rechallenge. The changes of weights were expressed as body weight loss prior to primary infection. (**C, D** and **E**) Detection of viral RNA in nasal swabs, throat swabs and anal swabs. SARS-CoV-2 RNA was detected by qRT-PCR in the swabs from seven monkeys at the indicated time points. (**F** and **G**) Hematological changes, including cell counts of WBC, LYMP and NEUT, as wells as the percentage of CD4^+^ T cells, CD8^+^ T cells and monocytes in peripheral blood were monitored respectively. (**H**) Levels of specific IgG against spike protein in each monkey. The levels of anti-viral antigen specific IgG from each monkey were detected at 3, 7, 14, 21, 28, 33 and 42 dpi. Four monkeys (M3-M6) were rechallenged at 28 dpi (the dotted line and shaded areas), and the results of initial infection and rechallenge were compared in bar graphs. The bars represented the average of four rechallenged animals at the indicated time points. The viral RNA in nasal, throat and anal swabs, of rechallenged animals were significantly lower than that of initial infection, while the specific antibodies were significantly increased. Meanwhile, significantly changes in hematological changes were observed between primary and second challenge (unpaired *t*-test, \**P<0.05, **P<0.01; ## P<0.01* 42 dpi vs 28 dpi).

**Figure 3.**
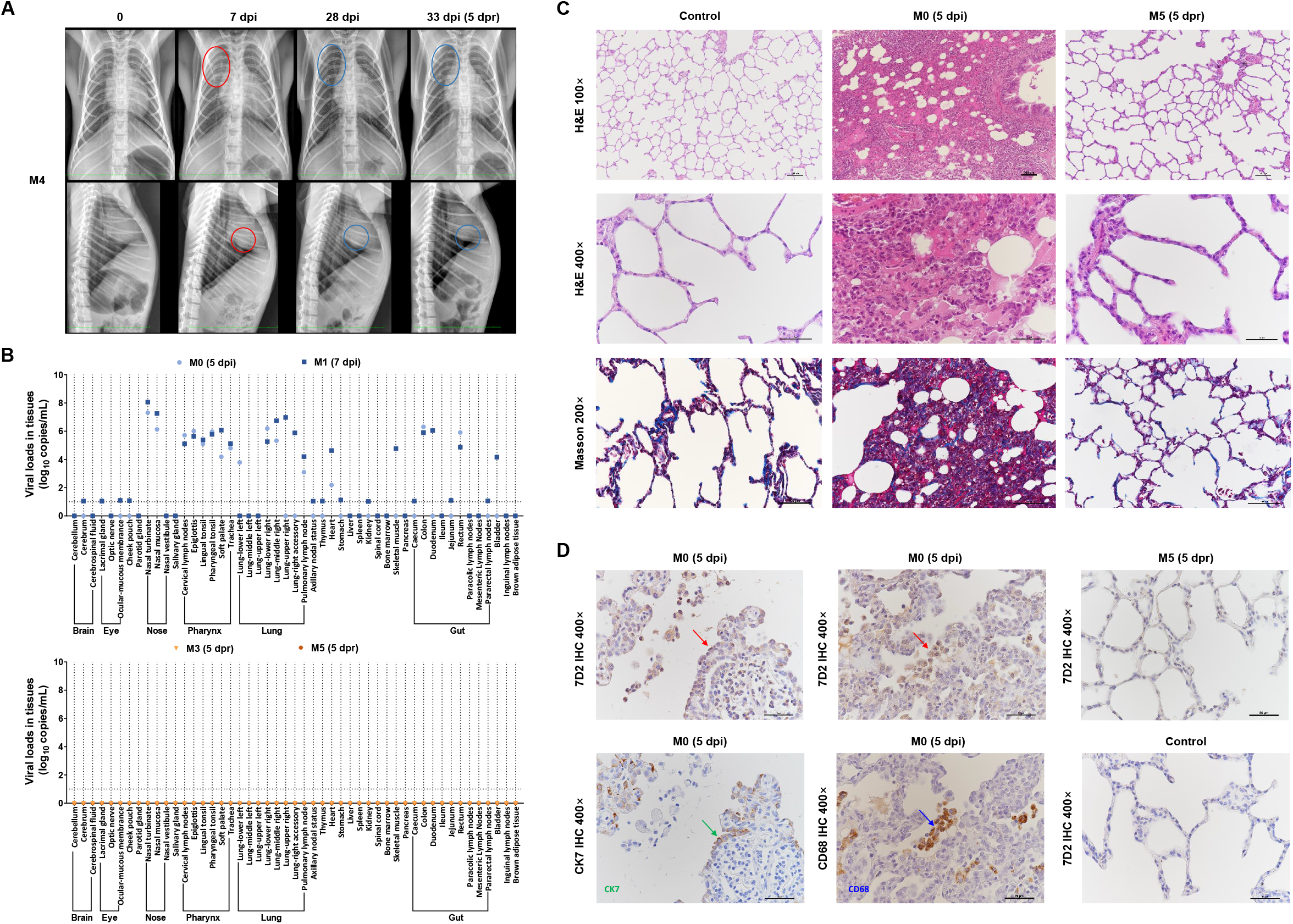
Comparison of Imaging, virus distribution and pathological changes between primary challenge stage and rechallenge stage. (**A**) Chest X ray of animals at 0, 7, 28 and 33 dpi (5 dpr) were examined and the photos of M4 was representatively shown. (**B**) Detection of viral RNA in the mainly organs, such as brain, eye, nose, pharynx, lung and gut. Compared to M0 and M1 with primary infection at 5 dpi or 7 dpi, viral replication tested negatively in the indicated tissues from M3 and M5 (at 5 dpr) with the virus rechallenge. Using viral load greater than 10 log10 copies/mL as threshold of positivity tissue-based PCR, tissues from 49 anatomical parts were detected for qualifying virus-infected positivity. 14 tissues from respiratory tract, gut and heart were shown SARS-CoV-2 positive cells from both M0 and M1. SARS-CoV-2 positive cells were only shown in left lower lung from M0 or in right upper lung, upper accessory lung, skeletal muscle, and bladder from M1 respectively. Remained tissues from 30 anatomical parts did not find SARS-CoV-2 positive cells, indicating these tissues were intact from viral invasion. (**C** and **D**) In M0 (5 dpi), an interstitial lesion including remarkedly widened alveolar septa and massive infiltrated inflammatory cells could be seen using HE staining. A mild fibrosis could be clearly seen within widened alveolar septa using Masson staining. Immunohistochemistry (IHC) against Spike protein of SARS-CoV-2 (7D2, red arrow), macrophage (CD68, blue arrow), or alveolar epithelial cell (CK7, green arrow) were in parallel visualized in Figure 3D. The Spike-positive cells overlapped with either alveolar epithelial cells or macrophages have shown the diffused interstitial pneumonia affected by SARS-CoV-2 invasion. In M5 (5 dpr), no remarked pathological changes and virus distribution were seen via HE staining, Masson staining or IHC, indicating the interstitial lesions have been completely recovered from SARS-CoV-2 primary infection and intact to reinfection. 100× or 200× Black scale bar = 100 μm. 400× Black scale bar = 50 μm. Data are representatives of three independent experiments.

Through HE staining, a mild to moderate interstitial pneumonia characterized by widened alveolar septa, increased alveolar macrophages and lymphocytes in the alveolar interstitium, and degenerated alveolar epithelia, and infiltrated inflammatory cells were shown in lung from monkeys with primary infection. Amount of collagen fiber could be also observed in the thickened alveolar interstitium in M0 and M1 monkeys by Modified Masson’s Trichrome stain at 5 or 7 dpi (Figure 3C). Meanwhile, the mucous membranes of trachea, tonsil, pulmonary lymph node, jejunum and colon in M0 and M1 exhibited inflammatory cell infiltrations (Figure S1), and infiltration with abundant CD4^+^ T cells, CD8^+^ T cells, B cells, macrophages and plasma cells in lung were specified by immunohistochemistry staining (IHC) (Figure S2). The virus-infected cells were mainly found in alveolar epithelia and macrophages by IHC on sequential sections (Figure 3D), as well as the mucous membranes of trachea, tonsil, pulmonary lymph node, jejunum and colon (Figure S1), confirming the SARS-CoV-2 could cause the COVID-19 in rhesus monkeys. Collectively, these data demonstrated that all the seven monkeys were successfully infected with SARS-CoV-2, and the characteristic of pathogenicity in monkey is similar to recent study(*9–13*).

At about 15 days posterior initial challenge, the body weight of infected monkeys (M2 to M6) gradually increased into normal range (4/5, except for M4, Figure 2A). All viral loads from nasopharyngeal and anal swabs returned negative (5/5, Figure 2C to 2E). In sera, spike protein-specific antibodies could be detected (5/5, Figure 2H, from 14 dpi). Chest X-ray resumed to normality at 28 dpi (5/5, Represented by M4, Figure 3A). These traits were similar to the discharging criteria including absence of clinical symptoms, radiological abnormalities and twice negative RT-PCR negativity in human infections (*14*). Taken together, it took about two weeks for monkeys undergone initial SARS-CoV-2 stepped into recovery stage (*10, 15*).

At 28 dpi, four monkeys (M3 to M6) undergone primary infection and recovery were rechallenged with the same dose of an identical SARS-CoV-2 strain intratracheally. The clinical tracking of the reinfection included weight loss (Figure 2A) and anal temperature (Figure 2B). A very interesting phenomenon was that the rechallenged monkeys exhibited a transiently increased temperature, which was not observed during the primary infection. Viral loads remained negative for a two-week intensive detection using nasopharyngeal and anal swabs post rechallenge of SARS-CoV-2 (Figure 2C to 2E). Peripheral blood measurements revealed no significant fluctuation during the rechallenging stage (Figure 2F and 2G). Moreover, no remarked abnormality of X-ray changes in M4 monkey at 33 dpi (5 dpr, Figure 3A). The only notable elevation was the concentration of antibodies against SARS-CoV-2 at 42 dpi (14 dpr), which is significantly higher than that at 28 dpi or 0 dpr (Figure 2H, ##*p*<0.01). Using necropsy specimens, there were no detectable viral RNA (Figure 3B, lower panel), significant pathological lesions (Figure 3C, Figure S1), virus-infected cells (Figure 3D, Figure S1) and immune cells infiltration (Figure S2) in lung and extrapulmonary tissue specimens from rechallenged monkeys (M3 and M5 at 5 dpr). Therefore, the rhesus monkeys with primary SARS-CoV-2 infection could not be reinfected with the identical strain during their early recovering stage.

To interpret the challenge-rechallenge disparity, it seemed to address valuable comparison of clinical, pathological and viral traits which comprehensively reflected the virus-host interaction between primary challenging stage and rechallenging stage in the four monkeys (M3 to M6). Firstly, viral loads from nasopharyngeal and anal swabs at 5 or 7 dpi were much higher than that at 5 or 7 dpr. Secondly, increased percentage of CD4^+^ T cells and decreased percentage of monocytes were observed at 7 dpr compared to that at 7 dpi. Thirdly, also of the most importance, the concentration of specific antibodies was much higher at 14 dpr than that at 14 dpi. The average titers of neutralizing antibodies exhibited linearly increased enhancement post primary infection (Table 1). Such gradually increased neutralizing antibodies against SARS-CoV-2 have provided an endurable humeral immunity aroused by primary infection, which might protect the same nonhuman primates from reinfection.

**Table 1.**
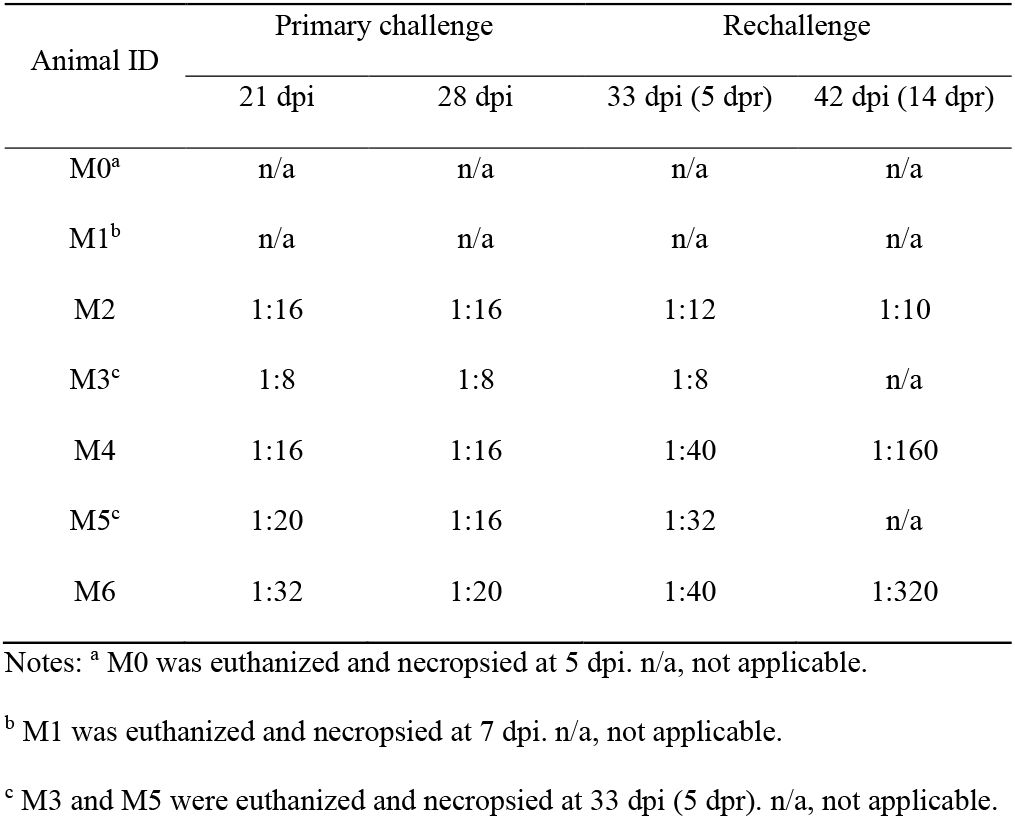
Neutralizing antibody titers to protect of SARS-CoV-2-infected Monkeys from reinfection.

| Animal ID | Primary challenge |  | Rechallenge |  |
| --- | --- | --- | --- | --- |
|  | 21 dpi | 28 dpi | 33 dpi (5 dpr) | 42 dpi (14 dpr) |
| M0<sup>a</sup> | n/a | n/a | n/a | n/a |
| M1<sup>b</sup> | n/a | n/a | n/a | n/a |
| M2 | 1:16 | 1:16 | 1:12 | 1:10 |
| M3<sup>c</sup> | 1:8 | 1:8 | 1:8 | n/a |
| M4 | 1:16 | 1:16 | 1:40 | 1:160 |
| M5<sup>c</sup> | 1:20 | 1:16 | 1:32 | n/a |
| M6 | 1:32 | 1:20 | 1:40 | 1:320 |

Notes: <sup>a</sup> M0 was euthanized and necropsied at 5 dpi. n/a, not applicable.
<sup>b</sup> M1 was euthanized and necropsied at 7 dpi. n/a, not applicable.
<sup>c</sup> M3 and M5 were euthanized and necropsied at 33 dpi (5 dpr). n/a, not applicable.

In the present challenge-rechallenge infection of SARS-CoV-2 in rhesus monkeys, observation and detection were within the relative short time window that neutralizing antibodies remained plateau after the primary infection. Moreover, all infected monkeys affected relative mild-to-moderate pneumonia, which is similar to mild or common clinical types of COVID-19 in the majority of the infected persons. Therefore, the immunity of primarily infected hosts which have been mildly impaired could be robustly resumed. Thirdly, mucosal immunity which have been aroused by primary infection including both respiratory and intestinal mucosal and local lymph nodes might contribute substantially against the newly attacked invasion of virus. A longer interval (longer than 6 months) between the primary challenge and re-challenge is needed to longitudinally track the host-virus interaction and elucidate the protective mechanism against SARS-CoV-2 in primates.

## Materials and Methods

### Ethics statement

Seven 3-to 5-year old rhesus macaques, named as M0 to M6, were housed and cared in an Association for the Assessment and Accreditation of Laboratory Animal Care (AAALAC)-accredited facility. All animal procedures and experiments were carried out in accordance with the protocols approved by the Institutional Animal Care and Use Committee (IACUC) of the Institute of Laboratory Animal Science, Chinese Academy of Medical Sciences (BLL20001). All animals were anesthetized with ketamine hydrochloride (10 mg/kg) prior to sample collection, and the experiments were performed in the animal biosafety level 3 (ABSL3) laboratory.

### Animal experiments

For primary infection, all animals were inoculated intratracheally with SARS-CoV-2 (SARS-CoV-2/WH-09/human/2020/CHN isolated in our laboratory) stock virus at a dosage of 10^6^ TCID50/1 mL inoculum volume. After the recovery, M3, M4, M5 and M6 were rechallenged intratracheally with the same dose (10^6^ TCID50/1 mL inoculum volume) SARS-CoV-2 at 28 dpi. To confirm the virus distribution and pathological changes, M0 at 5 dpi, M1 at 7 dpi, M3 and M5 at 33 dpi (5 dpr) were euthanasia and autopsied, respectively. All animals were monitored along the timeline to record body weights, body temperature, clinical signs, nasal/throat/anal swabs, hematological changes, immunocytes detection, chest X-ray and specific antibody. The animal experiment and longitudinal sampling schedule are shown in Figure 1.

### Quantification of SARS-CoV-2 RNA

The nasal/throat/anal swab samples and mainly tissue compartments collected from infected monkeys were tested for SARS-CoV-2 RNA by quantitative real-time reverse transcription-PCR (qRT-PCR). Total RNA was extracted and reverse transcription was performed as previously described (*16*). qRT-PCR reactions were carried out on an ABI 9700 Real-time PCR system (Applied Biosystems Instrument), the cycling protocol and the primers as follows: 50°C for 2 min, 95°C for 2 min, followed by 40 cycles at 95°C for 15 s and 60°C for 30 s, and then 95°C for 15 s, 60°C for 1 min, 95°C for 45 s. Forward primer: 5’-TCGTTTCGGAAGAGACAGGT-3’, Reverse primer: 5’-GCGCAGTAAGGATGGCTAGT-3’.

### Hematology

Whole blood was collected by EDTA-anticoagulation tube, and automatic hematology analyzer (ProCyte Dx) was used for hematological analysis. Hematologic parameters included the average total white blood cell counts (WBC), lymphocyte counts (LYPM), neutrophil counts (NEUT).

### Flow cytometry

Polychromatic flow cytometry was performed to analyze CD4^+^ T lymphocytes, CD8^+^ T lymphocytes and CD14^+^ monocytes. 50 μL of EDTA-anticoagulated whole blood were stained with the monoclonal antibodies CD3 BV605 (SP34-2, BD Biosciences, San Jose, CA), CD4 PerCP/Cyanine5.5 (Biolegend, 317428), CD8 FITC (Biolegend, 344704) and CD14 PE-Cy7 (Biolegend, 301814). The cells were resuspended in 1% paraformaldehyde and subjected to flow cytometry analysis within 24 hours. All the samples were analyzed by flow cytometry (FACSAria; BD, CA).

### ELISA

Sera were collected from each animal for the measurement of SARS-CoV-2 antibody by enzyme-linked immunosorbent assay (ELISA) along the detection timeline after the initial infection. 96-well plates were coated with 0.1 μg Spike protein of SARS-CoV-2 (Sino Biological, 40591-V08H) overnight at 4°C and blocked with 2% BSA/PBST for 1 hour at room temperature. 1:100 diluted sera were added to each well and incubated for 30 minutes at 37°C, followed by the HRP-labeled goat anti-monkey antibody (Abcam, ab112767) incubated for 30 minutes at room temperature. The reaction was developed by TMB substrate and determined at 450 nm.

### Histopathology and Immunohistochemistry

Autopsies were performed according to the standard protocol in ABSL3 laboratory at 5 or 7 dpi for M0 and M1, 5 dpr for M3 and M5. Tissues samples were fixed in 10% neutral-buffered formalin solution. Then, paraffin sections (3-4 μm in thickness) were prepared and stained with Hematoxylin and Eosin (H&E) and modified Masson’s Trichrome stain (Masson) prior to the observation by light microscopy. For immunohistochemistry (IHC) staining to identify the cell type and the expression of SARS-CoV-2 antigen, paraffin dehydrated sections (3-4 μm in thickness) were treated with an antigen retrieval kit (Boster, AR0022) for 1 min at 37°C and quenched for endogenous peroxidases in 3% H2O2 in methanol for 10 min. After blocking in 1% normal goat serum for 1 hour at room temperature, the sections were stained with 7D2 monoclonal antibody (1:500 dilution, laboratory preparation) and CD68 antibody (1:500 dilution, Abcam, ab201340) at 4°C overnight, following with the incubation of HRP-labeled goat anti-mouse IgG (Beijing ZSGB Biotechnology, ZDR-5307) for 1 hour. Alternatively, the sections were stained with CK7 antibody (1:1000 dilution, Abcam, ab181598), CD4 antibody (1:500 dilution, Beijing ZSGB Biotechnology), CD8 antibody (1:200 dilution, Abcam, ab4055), CD20 antibody (1:500 dilution, Abcam, ab78237) or CD138 antibody (1:500 dilution, Abcam, ab128936) at 4°C overnight, followed by HRP-labeled goat anti-rabbit IgG secondary antibody (Beijing ZSGB Biotechnology, PV9001) for 60 min. Then, the sections were visualized by incubation with 3,30-diaminobenzidine tetrahydrochloride (DAB) and the image was viewed under an Olympus microscope. The tissue sections (stored by our laboratory) of DMEM (1 mL inoculation volume by intratracheal route)-treated monkey were used as a negative control.

### Neutralizing antibody assay

Sera samples were tested for the presence of neutralizing antibody observed by cytopathic effect (CPE). Briefly, the sera from monkeys were heat-inactivated at 56 °C for 30 min. Then, serially two-fold diluted sera were incubated with 100 TCID50 SARS-CoV-2 for 1 h at 37°C, and added into Vero-E6 cells in a 96-well-plate. Cells were cultured for 1 week to observe for CPE and the serum dilution in which 50% of the cells were protected from infection was calculated. Each dilution of serum was tested in triplicates.

### Statistical analysis

Comparisons between the two groups were determined using two-tailed unpaired Student’s or Welch’s *t*-test. All data were analyzed with GraphPad Prism 8.0 software. The level of statistical significance is designated as \**p* < 0.05, \*\**p* < 0.01.

## ACKNOWLEDGEMENTS

This work was supported by the CAMS initiative for Innovative Medicine of China (Grant No. 2016-I2M-2-006), National Mega projects of China for Major Infectious Diseases (Grant No. 2017ZX10304402) and National Key Research and Development Project of China (Grant No. 2016YFD0500304).

## AUTHOR CONTRIBUTIONS

Conceptualization: C.Q.; Methodology: L.B., W.D., H.G., C.X., J.L., J.X. and Q.L.; Investigation: L.B., W.D., H.G., C.X., J.L., J.X., Q.L., J.L., P.Y., Y.X., F.Q., Y.Q., F.L., Z.X., H.Y., S.G., M.L., G.W., S.W., Z.S., Y.L., W.Z., Y.H., L.Z., X.L. and Q.W.; Writing – Original Draft: J.X.; Writing –Review and Editing: J.X. and C.Q.; Funding Acquisition: L.B. and C.Q.; Resources: C.Q.; Supervision: C.Q.

---

**Supplemental Figure 1.**
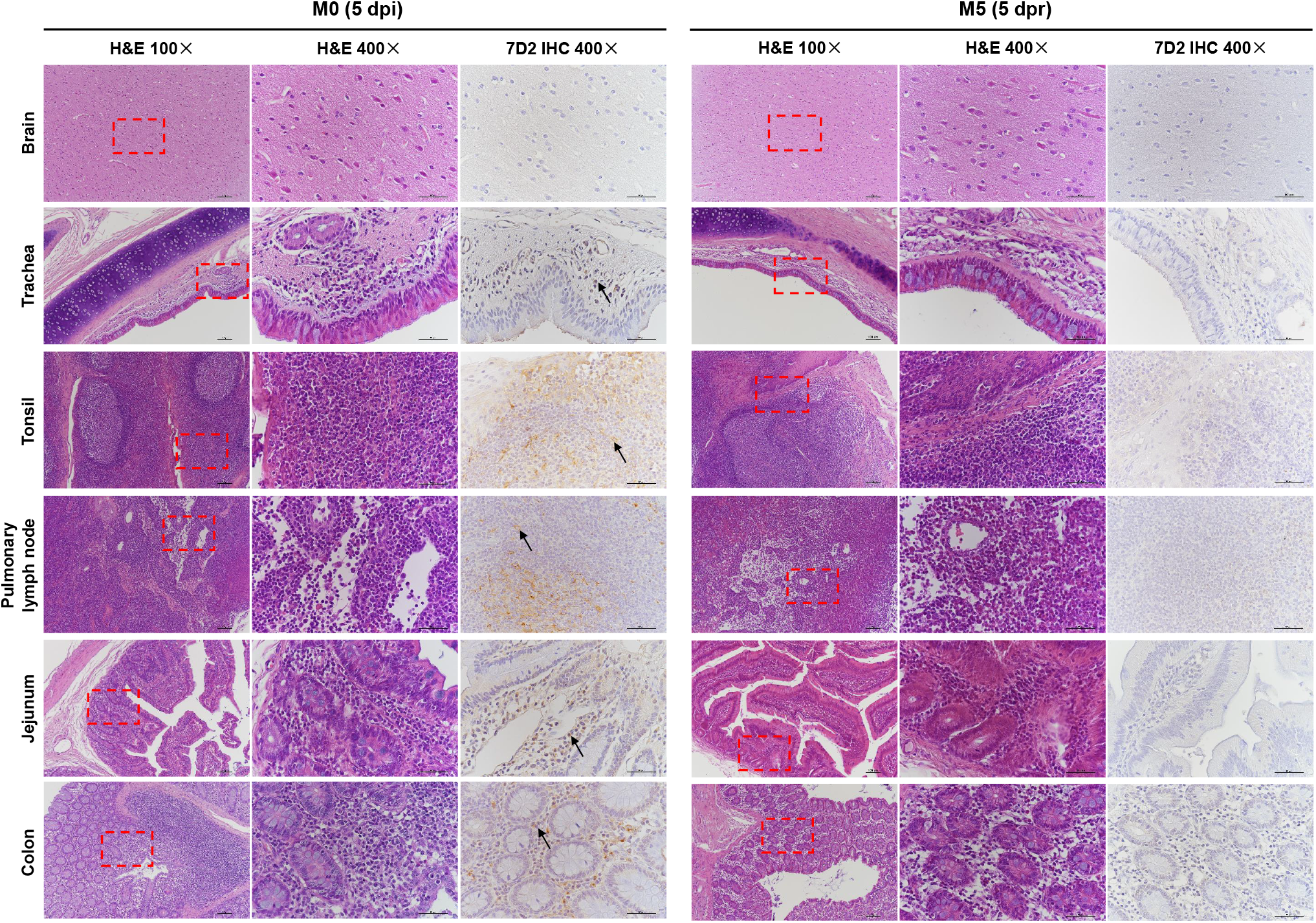
Comparison of virus distribution and pathological changes in extrapulmonary organs between primary challenge stage and rechallenge stage. The pathological changes were observed by HE staining, and the viral antigens were detected by Immunohistochemistry (IHC) against Spike protein of SARS-CoV-2 (7D2, black arrow). In M0 (5 dpi), inflammatory cell infiltrations and viral antigens were observed in mucous membranes of trachea, tonsil, pulmonary lymph node, jejunum and colon, and no lesions were observed in brain. Compared to M0, no remarked pathological changes and virus distribution were seen via HE staining or IHC in M5 (5 dpr), indicating the animal have been completely protected from rechallenge. The red frames are the area of magnification. 100× Black scale bar = 100 μm. 400× Black scale bar = 50 μm. Data are representatives of three independent experiments.

**Supplemental Figure 2.**
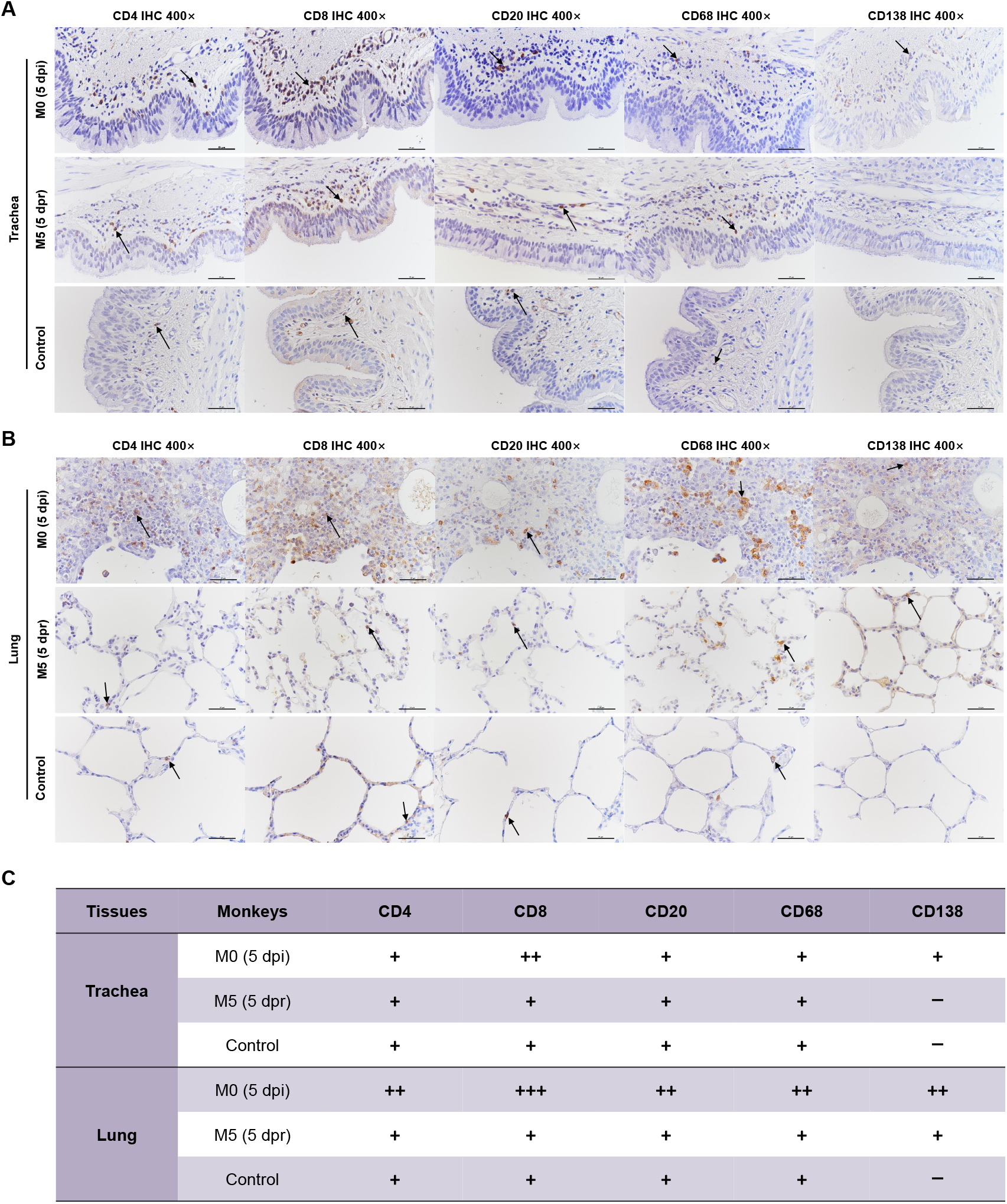
Comparison of immune cells distribution in trachea and lung between primary challenge stage and rechallenge stage. The immune cells in trachea (**A**) and lung (**B**) were detected by Immunohistochemistry (IHC) respectively (CD4 and CD8 for T cell, CD20 for B cell, CD68 for macrophage and CD138 for plasma cells), and then the number of positive cells were evaluated and scored (**C**). Compared to control, the lung of M0 (5 dpr) was infiltrated with plenty of CD4^+^ T cell, CD8^+^ T cell, B cells, macrophages and plasma cells, while abundant CD8^+^ T cell and scattered plasma cells were also observed in trachea. No obvious difference in immune cells distribution was observed in M5 (5 dpr) compared to control except the increased plasma cells in lung. 400× Black scale bar = 50 μm. Data are representatives of three independent experiments.

## REFERENCE

1. N. Zhu et al., A Novel Coronavirus from Patients with Pneumonia in China, 2019. N Engl J Med, 10.1056/NEJMoa2001017 (2020).

2. Coronavirus disease 2019 (COVID-19) Situation Report. WHO, (2020). At https://www.who.int/emergencies/diseases/novel-coronavirus-2019/situation-reports/.

3. L. Lan et al., Positive RT-PCR Test Results in Patients Recovered From COVID-19. Jama 323, 1502–1503 (2020).

4. L. Zhou, K. Liu, H. G. Liu, Cause analysis and treatment strategies of “recurrence” with novel coronavirus pneumonia (covid-19) patients after discharge from hospital. Zhonghua jie he he hu xi za zhi = Zhonghua jiehe he huxi zazhi = Chinese journal of tuberculosis and respiratory diseases 43, E028 (2020).

5. J. An et al., Clinical characteristics of the recovered COVID-19 patients with re-detectable positive RNA test. medRxiv, 2020.2003.2026.20044222 (2020).

6. F. Wu et al., Neutralizing antibody responses to SARS-CoV-2 in a COVID-19 recovered patient cohort and their implications. medRxiv, 2020.2003.2030.20047365 (2020).

7. N. M. A. Okba et al., SARS-CoV-2 specific antibody responses in COVID-19 patients. medRxiv, 2020.2003.2018.20038059 (2020).

8. K. K. To et al., Temporal profiles of viral load in posterior oropharyngeal saliva samples and serum antibody responses during infection by SARS-CoV-2: an observational cohort study. The Lancet. Infectious diseases, (2020).

9. S. Lu et al., Comparison of SARS-CoV-2 infections among 3 species of non-human primates. bioRxiv, 2020.2004.2008.031807 (2020).

10. V. J. Munster et al., Respiratory disease and virus shedding in rhesus macaques inoculated with SARS-CoV-2. bioRxiv, 2020.2003.2021.001628 (2020).

11. B. Rockx et al., Comparative pathogenesis of COVID-19, MERS, and SARS in a nonhuman primate model. Science, (2020).

12. B. N. Williamson et al., Clinical benefit of remdesivir in rhesus macaques infected with SARS-CoV-2. bioRxiv, 2020.2004.2015.043166 (2020).

13. P Yu et al., Age-related rhesus macaque models of COVID-19. Animal models and experimental medicine 3, 93–97 (2020).

14. Diagnostic and treatment protocol for Novel Coronavirus Pneumonia (Trial version 6). General Office of National Health Commission, (2020). At http://www.nhc.gov.cn/yzygj/s7652m/202002/54e1ad5c2aac45c19eb541799bf637e9.shtml.

15. S. F. Wang et al., Antibody-dependent SARS coronavirus infection is mediated by antibodies against spike proteins. Biochemical and biophysical research communications 451, 208–214 (2014).

16. L. Bao et al., The Pathogenicity of SARS-CoV-2 in hACE2 Transgenic Mice. bioRxiv, 2020.2002.2007.939389 (2020).

